# UCell: robust and scalable single-cell gene signature scoring

**DOI:** 10.1101/2021.04.13.439670

**Authors:** Massimo Andreatta, Santiago J. Carmona

## Abstract

UCell is an R package for evaluating gene signatures in single-cell datasets. UCell signature scores, based on the Mann-Whitney U statistic, are robust to dataset size and heterogeneity, and their calculation demands less computing time and memory than other available methods, enabling the processing of large datasets in a few minutes even on machines with limited computing power. UCell can be applied to any single-cell data matrix, and includes functions to directly interact with Seurat objects. The UCell package and documentation are available on GitHub at https://github.com/carmonalab/UCell

## 1 Introduction

In single-cell RNA-seq analysis, gene signature (or “module”) scoring offers a versatile approach for the identification of cell types, states and active biological processes. The Seurat R package (Stuart *et al*., 2019) is one of the most comprehensive and widely used frameworks for scRNA-seq data analysis. Seurat provides a computationally efficient gene signature scoring function, named AddModuleScore, originally proposed by (Tirosh *et al*., 2016). However, because genes are binned based on their average expression across the whole dataset for normalization purposes, the method generates inconsistent results for the same cell depending on the composition of the dataset. Inspired by the AUCell algorithm implemented in SCENIC (Aibar *et al*., 2017), we propose UCell, a gene signature scoring method based on the Mann-Whitney U statistic. UCell scores depend only on the relative gene expression in individual cells and are therefore not affected by dataset composition. We provide a time-and memory-efficient implementation of the algorithm that can be seamlessly incorporated into Seurat workflows.

## 2 Methods

UCell calculates gene signature scores for scRNA-seq data based on the Mann-Whitney U statistic (Mann and Whitney, 1947). Given a *m × n* matrix *M* of numerical values (e.g. gene expression measurements) for *m* genes in *n* cells, we first calculate the relative ranks *r*_*m,n*_ of the scores in each column *n*; in other words, a ranked list of genes for each cell in the dataset. Because of the well-known drop-out effect, single-cell data commonly contain many zero measurements, resulting in a long tail of bottom-ranking genes. To mitigate this uninformative tail, we set *r*_*m,n*_ = *r*_*max*_ +1 for all *r*_*m,n*_ > *r*_*max*_, with *r*_*max*_ = 1500 by default (matching typical thresholds used for quality control for minimum number of genes detected). To evaluate a gene signature *G* composed of *y* genes *g*_*1*_, …,*g*_*y*_ we calculate a score *U*_*C*_ for each cell *n* in *M* with the formula:

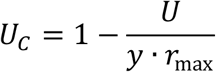

where U is the Mann-Whitney U statistic calculated by:

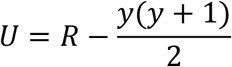

and R is the sum of the relative ranks *r*_*{i in G},n*_ of the genes in signature G.

We note that U score is closely related to the area-under-the-curve (AUC) statistic for ROC curves (Mason and Graham, 2002), therefore we expect UCell scores to correlate with methods based on AUC scores such as AUCell (Aibar *et al*., 2017). Internally, UCell uses the *frank* function from the *data*.*table* package for efficient ranks computations. Large datasets are automatically split into chunks of reduced size, which can be processed serially (minimizing memory usage) or in parallel through the *future* package (Bengtsson, 2020) (minimizing execution time) depending on the available computational resources.

## 3 Results

UCell is an R package for the evaluation of gene signature enrichment designed for scRNA-seq data. Given a gene expression matrix or Seurat object, and a list of gene sets, UCell calculates signature scores for each cell, for each gene set. In the following illustrative example, we applied UCell to a single-cell multimodal dataset of human blood T cells (Hao *et al*., 2020), which were annotated by the authors using both gene (scRNA-seq) and cell surface marker expression (CITE-seq) (Figure 1A). Provided a set of T cell subtype-specific genes (Table 1), UCell helps interpreting clusters in terms of signature enrichment in low-dimensional spaces such as the UMAP (Figure 1B). Importantly, UCell scores are based on the relative ranking of genes for individual cells, therefore they are robust to dataset composition. Evaluating a CD8 T cell signature on the full dataset or on CD8 T cells only, results in identical score distributions for CD8 T cells in the two settings (Figure 1C). Conversely, AddModuleScore from Seurat normalizes its scores against the average expression of a control set of genes across the whole dataset, and is therefore dependent on dataset composition. CD8 T cells analyzed in isolation or in the context of the full T cell dataset are assigned highly different AddModuleScore scores, with median∼1 in the full dataset and median∼0 for the CD8 T cell subset (Figure 1D). Another widely-used method for single-cell signature scoring, AUCell (Aibar *et al*., 2017), is also based on relative rankings and therefore has the same desirable property as UCell of reporting consistent scores regardless of dataset composition. Compared to AUCell, UCell is about three times faster (Figure 1E) and uses significantly less memory (Figure 1F). For example, AUCell requires over 64GB of RAM to process 100,000 single-cells, while UCell uses only 5.5GB of peak memory, making it suitable even for machines with limited computing power.

**Table 1:** Gene signatures for T cell subsets in Figure 1

| T cell type | Gene set |
| --- | --- |
| CD4 T cell | *CD4*, *CD40LG* |
| CD8 T cell | *CD8A*, *CD8B* |
| Treg | *FOXP3*, *IL2RA* |
| MAIT | *KLRB1*, *SLC4A10*, *NCR3* |
| gd T cell | *TRDC*, *TRGC1*, *TRGC2*, *TRDV1* |

**Figure 1:**
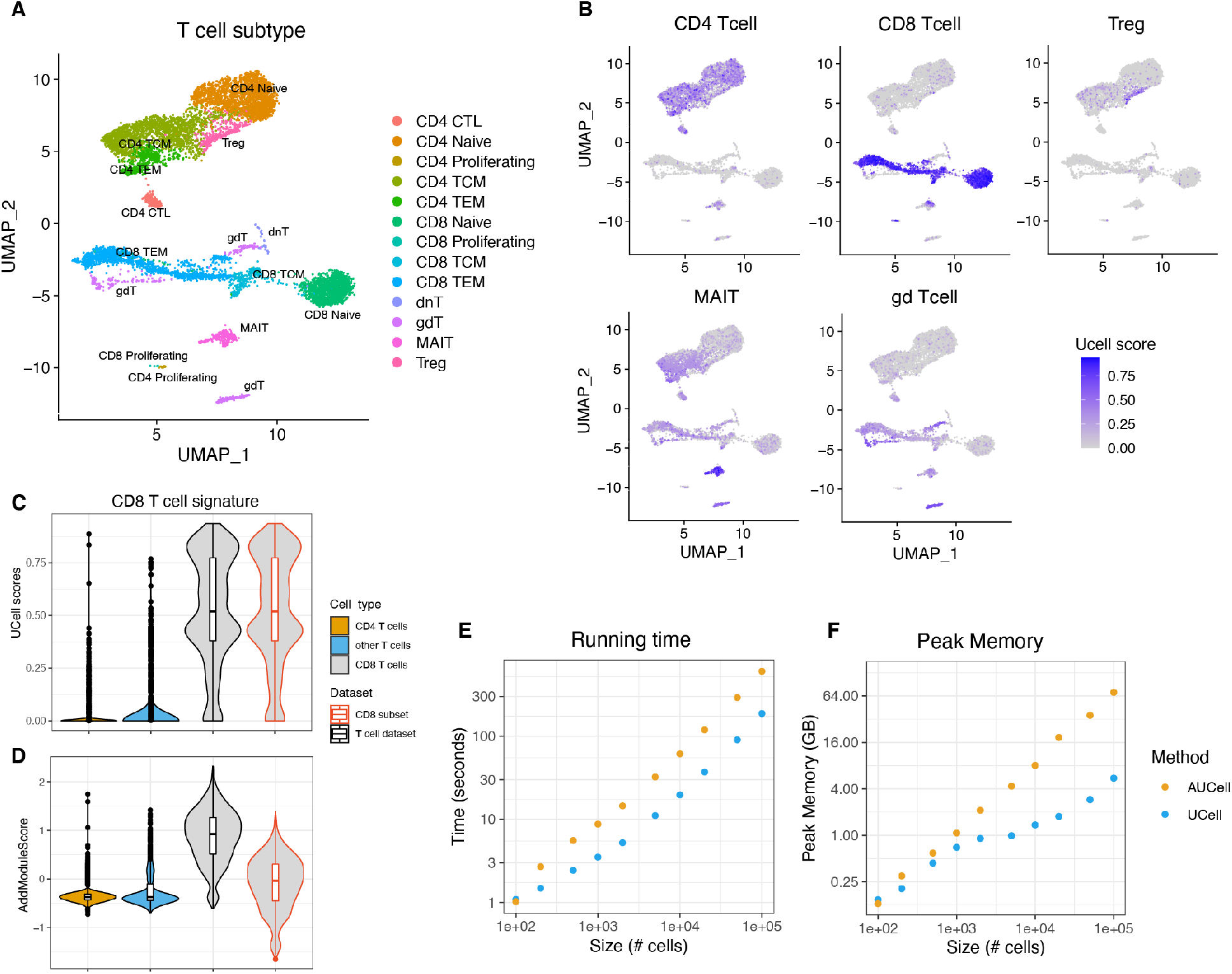
Evaluating T cell signatures using UCell. A) UMAP representation of T subsets from the single-cell dataset by (Hao *et al*., 2020). **B)** UCell score distribution in UMAP space for five gene signatures evaluated using UCell. **C-D)** Comparison of UCell score **(C)** and Seurat’s AddModuleScore **(D)** distributions for a two-gene CD8 T cell signature (*CD8A, CD8B*), evaluated on the complete T cell dataset (black outlines), or on the subset of CD8 T cells only (red outlines); UCell scores for CD8 T cell have the same distribution in the complete or subset dataset, while AddModuleScores are highly dependent on dataset composition. **E-F)** Running time **(E)** and peak memory **(F)** for UCell and AUCell (which produces similar results) on datasets of different sizes show that UCell is about three times faster and requires up to ten times less memory on large (>10^4^) single-cell datasets.

The code to generate these results and figures is available at https://gitlab.unil.ch/carmona/UCell_demotogether with further UCell applications. The UCell package and documentation are available on GitHub at https://github.com/carmonalab/UCell

## References

Aibar,S. et al. (2017) SCENIC: single-cell regulatory network inference and clustering. Nat. Methods, 14, 1083–1086.

Bengtsson,H. (2020) A Unifying Framework for Parallel and Distributed Processing in R using Futures. ArXiv200800553 Cs Stat.

Hao,Y. et al. (2020) Integrated analysis of multimodal single-cell data Genomics.

Mann,H.B. and Whitney,D.R. (1947) On a Test of Whether one of Two Random Variables is Stochastically Larger than the Other. Ann. Math. Stat., 18, 50–60.

Mason,S.J. and Graham,N.E. (2002) Areas beneath the relative operating characteristics (ROC) and relative operating levels (ROL) curves: Statistical significance and interpretation. Q. J. R. Meteorol. Soc., 128, 2145–2166.

Stuart,T. et al. (2019) Comprehensive Integration of Single-Cell Data. Cell, 177, 1888-1902.e21.

Tirosh,I. et al. (2016) Dissecting the multicellular ecosystem of metastatic melanoma by single-cell RNA-seq. Science, 352, 189–196.

